# Gene Expression Prediction from Histology Images via Hypergraph Neural Networks

**DOI:** 10.1101/2024.08.05.606608

**Authors:** Bo Li, Yong Zhang, Qing Wang, Chengyang Zhang, Mengran Li, Guangyu Wang, Qianqian Song

**Author notes:** Corresponding authors: Qianqian Song, PhD; Yong Zhang, PhD. **Biographical Note** **Bo Li** His research focuses on biomedical image analysis and bioinformatics. **Yong Zhang** Faculty of Information Technology, Beijing Institute of Artificial Intelligence, Beijing University of Technology, Beijing, China. His research focuses on big data analysis and visualization, and the applications of artificial intelligence. **Qing Wang** Key Laboratory of Intelligent Education Technology and Application of Zhejiang Province, Jinhua, China. His research interests include data mining, artificial intelligence, and their applications. **Chengyang Zhang** His research focuses on computer vision and deep generative model. **Mengran Li** School of Intelligent Systems Engineering, Sun Yat-sen University, Guangzhou, Shenzhen, China. His research interests include big data and artificial intelligence. **Guangyu Wang** Houston Methodist Research Institute, Weill Cornell Medical College. His research focuses on developing computational models for analyzing high-throughput multi-omics datasets and interrogating how cell fate is regulated. **Qianqian Song** Her research focuses on developing innovative computational methods to decipher disease mechanisms and identify therapeutic biomarkers.

## Abstract

Spatial transcriptomics reveals the spatial distribution of genes in complex tissues, providing crucial insights into biological processes, disease mechanisms, and drug development. The prediction of gene expression based on cost-effective histology images is a promising yet challenging field of research. Existing methods for gene prediction from histology images exhibit two major limitations. First, they ignore the intricate relationship between cell morphological information and gene expression. Second, these methods do not fully utilize the different latent stages of features extracted from the images. To address these limitations, we propose a novel hypergraph neural network model, HGGEP, to predict gene expressions from histology images. HGGEP includes a gradient enhancement module to enhance the model’s perception of cell morphological information. A lightweight backbone network extracts multiple latent stage features from the image, followed by attention mechanisms to refine the representation of features at each latent stage and capture their relations with nearby features. To explore higher-order associations among multiple latent stage features, we stack them and feed into the hypergraph to establish associations among features at different scales. Experimental results on multiple datasets from disease samples including cancers and tumor disease, demonstrate the superior performance of our HGGEP model than existing methods.

**Key Points:** We develop a novel histology image-based gene prediction model named HGGEP, which demonstrates high accuracy and robust performance.

To reveal the intricate relationship between cell morphology and gene expression in images, we propose a gradient enhancement module, which effectively improves the model’s capability in perceiving cell morphology in images.

HGGEP includes a hypergraph module that efficiently models higher-order associations among features across multiple latent stages, resulting in significant performance improvement.

## INTRODUCTION

Diverse cell types are intricately arranged both spatially and structurally within tissues to fulfill their specific functions. Revealing the intricate spatial architecture and cell activities within heterogeneous tissues holds considerable importance in comprehending the cellular mechanisms and functions associated with diseases^1-4^. Spatial Transcriptomics (ST) serves as an advanced technology that can be utilized to elucidate the spatial distribution of genes at both tissue and spot levels. This technology has significantly advanced our understanding of gene expressions in biological processes^5^, playing a crucial role in exploring disease mechanisms^6^ and revealing novel drug targets^7^. The rapid progress of ST technology allows for the simultaneous analysis of gene expression, cell or spot locations, and corresponding histology images. Currently, numerous researchers are actively engaged in related studies, covering spatial domain recognition ^8-10^, spatial transcriptomics deconvolution^11-13^, and inference of spatial cellular interactions^14, 15^.

However, the considerable cost associated with acquiring spatial transcriptomics data limits the widespread pursuit of research on ST technologies. In contrast, histology images of various disease tissues are more accessible. Recently, researchers have shifted their focus toward predicting gene expression from whole-slide image (WSI) data. Some methods, such as ST-Net^16^, HisToGene^17^, Hist2ST^18^, DeepPT^19^, BLEEP^20^, and THItoGene^21^, have emerged for this purpose. Initially, ST-Net pioneers the use of deep learning techniques to predict spatial gene expression from WSI, yielding promising results. HisToGene and Hist2ST improve the prediction performance by incorporating transformer models to capture global associations of image features across different spots in a WSI. Meanwhile, Hist2ST leverages graph neural networks^22^ to enhance local associations of image features between spots. Recently, THItoGene further employs graph attention networks to explore the correlation between gene expression and spatial location. Unlike the direct prediction methods from images to gene expression mentioned above, BLEEP uses a contrastive learning approach to align images with gene expression. It’s worth mentioning that Adam et al.^23^conduct a comprehensive benchmark in the current field. They provide multiple metrics to thoroughly evaluate various models, including the performance of predicted gene expression, model generalizability, translational potential, usability, and computational efficiency of each method.

Although the aforementioned work has made significant progress, several important aspects remain overlooked. For instance, ST-Net ignores the positional information of spots and does not explore the correlations between multiple spots. HisToGene directly feeds image patches into ViT, resulting in significant information loss. Hist2ST and THItoGene use graph neural networks and graph attention networks, respectively, to model spot-neighborhood relations within global features, but they overlook the associations between spots that are spatially distant yet closely related. Overall, these existing methods still face two primary limitations: (1) they ignore the intricate relationship between cell morphological information and gene expression; (2) insufficient utilization of image-based features at multiple latent stages, coupled with the oversight of high-order associations among those features. A detailed comparison of existing methods is shown in **Supplementary Table 1**.

Regarding the first limitation, existing methods based on traditional convolution primarily concentrate on semantic information, i.e., pixel values in the image. They do not sufficiently consider the gradient relationship between the current position and its neighboring positions, which leads to the difficulty of the model to perceive the cell morphological information related to gene expression. To address this limitation, our HGGEP model includes the gradient enhancement module to refine the extracted imaging features and generate latent feature maps with prominent cell morphological information. To address the second limitation and enhance the utilization of features at multiple latent stages within WSI, our HGGEP model employs a two-step strategy. Specifically, HGGEP first extracts multiple latent features from WSI through a lightweight backbone network^24^, and subsequently refine the representation of features at each latent stage using the attention mechanism^25, 26^. To explore higher-order associations among multiple latent stage features, we innovatively introduce a hypergraph association module based on multiple metrics. Compared to traditional graph neural networks^27^, hypergraph convolution^28, 29^ can multiple nodes with a single hyperedge, enabling effective joint representation and modeling of higher-order associations among features. Collectively, we propose a novel HGGEP model that overcomes existing challenges and achieves superior performance in gene expression prediction from histology images.

## RESULTS

### Overview of HGGEP model

Our hypergraph neural network-based model, HGGEP (HyperGraph Gene Expression Prediction), is depicted in **Figure 1**. The process initiates with the partitioning of a whole slide image into multiple patches centered around spots. Acknowledging the intricate relationship between cell morphology and gene expression, the Gradient Enhancement Module (GEM) is utilized to enhance the model’s perception of cell morphology. The latent features preprocessed by GEM undergo processing through a lightweight backbone network^24^ to extract features at multiple latent stages. These latent features are then fed into the Convolutional Block Attention Module (CBAM^25^) and Vision Transformer (ViT^26^), utilizing attention mechanisms to optimize the representation of features at each latent stage. To uncover high-order associations among features across different stages, a Hypergraph Association Module (HAM) based on nearby positions and distance relations is employed for global associations modeling and local representation. Following this, the latent features output from the ViT and HAM are concatenated and fed into the Long Short-Term Memory (LSTM^30, 31^) module, enhancing information exchange among latent features. Ultimately, the latent features from the last layer are input into a gene regression head to yield the predicted gene expression on this histology image.

**Figure 1.**
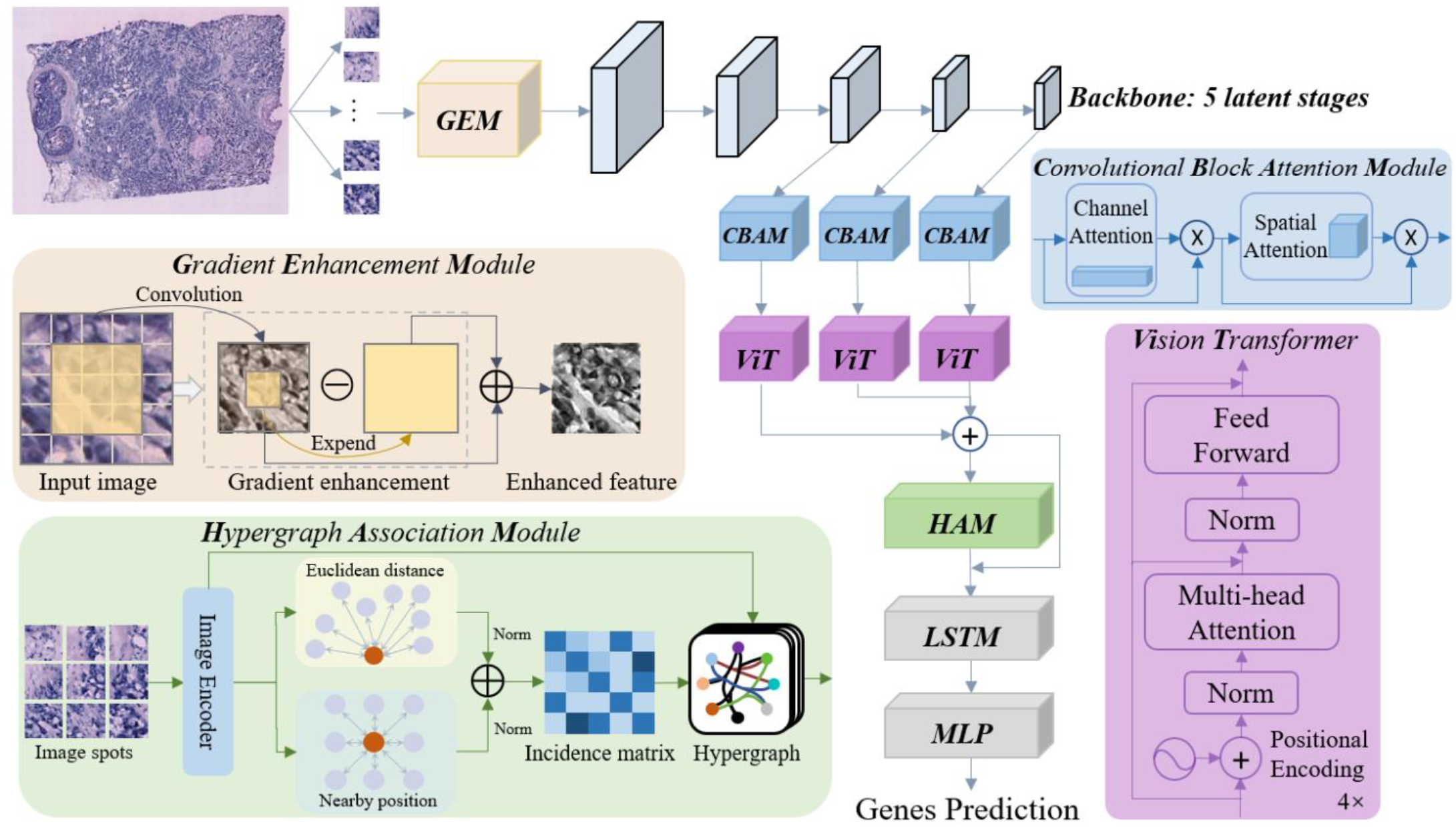
Overview of our HGGEP model. This model consists of three pivotal components: GEM, designed to capture the intricate relationship between cell morphology and gene expression; CBAM and Vision Transformer modules, employed for extracting internal features at each latent stage; and a Hypergraph Association Module (HAM), dedicated to revealing higher-order associations among multiple latent stage features.

### Performance evaluation using diverse spatial transcriptomics datasets

To evaluate the performance of HGGEP, we conducted experiments using a leave-one-out validation approach on a total of 44 sections from human HER2-positive breast tumor (HER2+^32^) dataset (section A1-G3) and human cutaneous squamous cell carcinoma (cSCC^33^) dataset (section P2_ST_rep1-P10_ST_rep3). The performance of HGGEP is compared with existing methods including Hist2ST, HisToGene, and ST-Net. **Figure 2** illustrates the comparison results based on the Pearson Correlation Coefficient (PCC) index for each method. Specifically, on the HER2+ datasets, the HGGEP model exhibits an average PCC index and median PCC index approximately 5% higher than the second-best-performing Hist2ST model. Similarly, on the cSCC datasets, the HGGEP model outperforms the Hist2ST model by around 4%. **Figure 2** reveals that most of the compared models perform less favorably in sections A2-A6 and F1-F3, with PCC indices consistently below 0.07. In contrast, our HGGEP model successfully improves the performance by approximately 5%. Notably, the most substantial improvement relative to the compared models occurs in sections C1-C5, with the PCC increase of about 7%. These results demonstrate that compared with existing methods, our HGGEP model presents superior capabilities of predicting gene expressions from histology images. Additionally, we conduct experiments to evaluate the model’s goodness-of-fit on the training dataset, as shown in Figure S4 of the supplementary materials. The results indicate that the proposed HGGEP model demonstrates excellent goodness-of-fit on the training dataset and significantly outperforms other models in overall performance.

**Figure 2.**
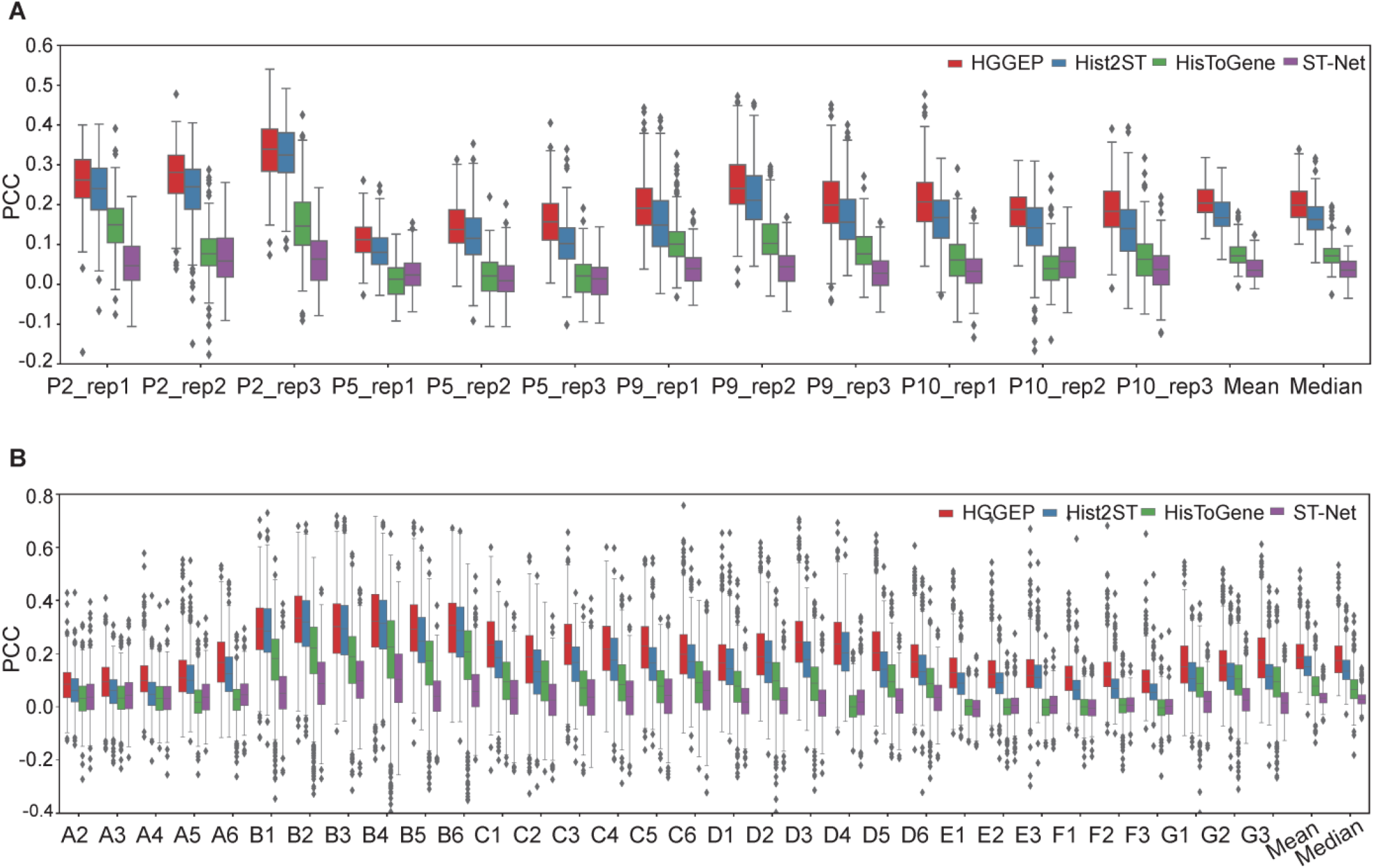
Benchmark of the gene expression prediction performance. Comparison results between our HGGEP model and existing methods on the (A) HER2+ datasets and (B) and cSCC datasets.

To comprehensively validate the gene expression prediction performance of our model, we also provide performance comparisons using other metrics, including Structural Similarity Index Measure (SSIM) and Root Mean Square Error (RMSE). SSIM helps evaluate the model’s ability to preserve the spatial patterns and structural features of gene expression. RMSE quantifies the difference between the predicted gene expression values and the actual observations by calculating the square root of the average squared prediction errors. Therefore, a higher SSIM and a lower RMSE indicate better model performance. **Figure 3** presents the comparison results of various models based on SSIM and RMSE metrics. Clearly, HGGEP achieves the highest SSIM value and the lowest RMSE value, demonstrating that the HGGEP model proposed in this paper achieves the best gene expression prediction performance on these two metrics. Additionally, we included comparisons with THItoGene in the supplementary file (Figure S1-S3). The results still indicate that our method achieved the best performance.

**Figure 3.**
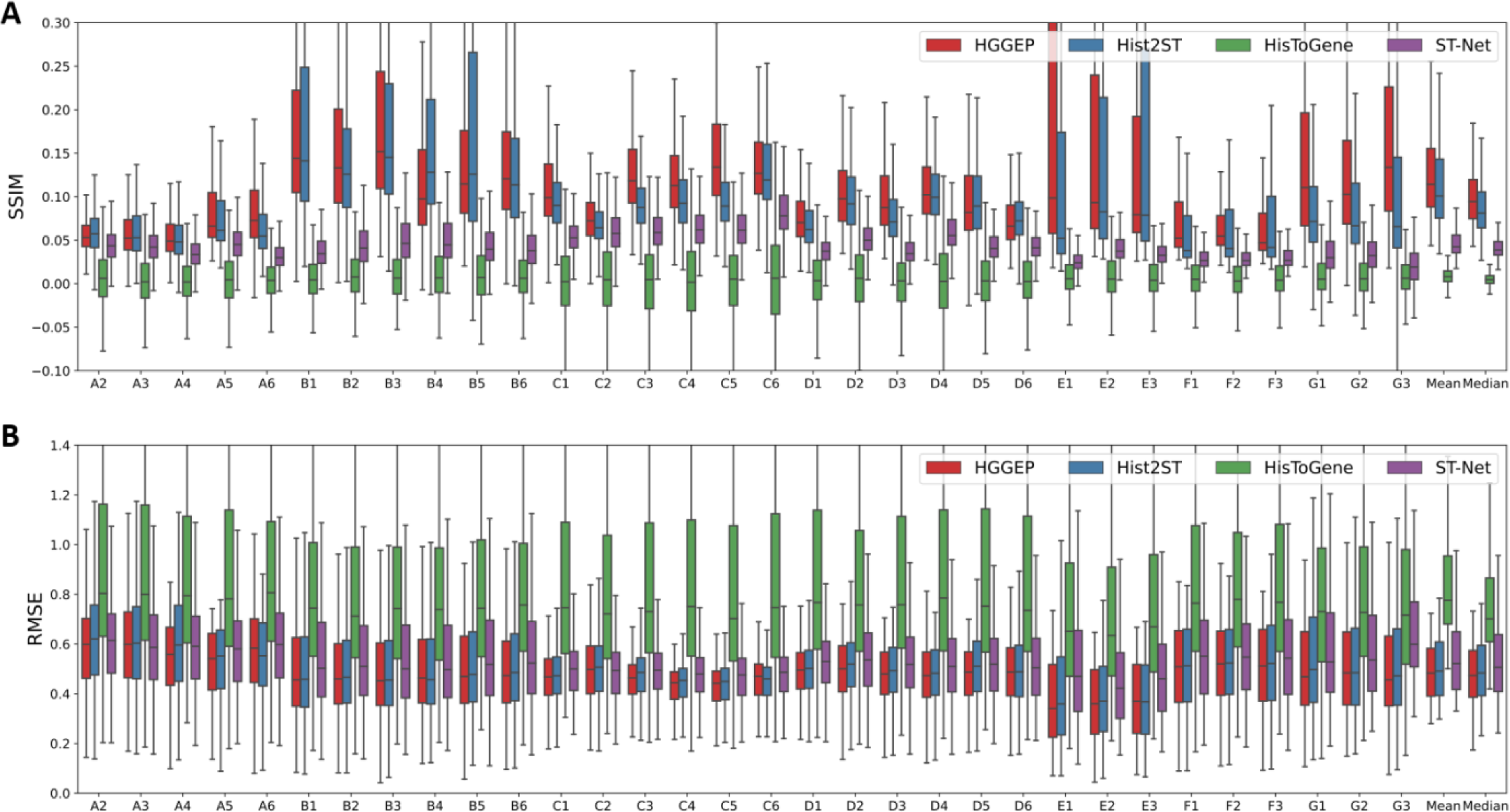
Benchmark of the gene expression prediction performance based on SSIM and RMSE. Comparison results between our HGGEP model and existing methods for SSIM (A) and RMSE (B) on the HER2+ datasets. Among them, a higher SSIM and a lower RMSE indicate better model performance.

To assess the contribution of each module’s performance, ablation experiments were conducted on the HER2+ datasets, and the results are presented in **Figure 4**. In this figure, red color represents the performance of ablation experiments conducted on the HGGEP model, while the other three colors correspond to the compared methods (Hist2ST, HisToGene, ST-Net). The observations in **Figure 4** lead to the following three conclusions: 1) As the number of modules removes, the mean PCC of the model gradually declines, underscoring the substantial contribution each module makes to the overall performance of the HGGEP model. 2) The results from the ablation experiments indicate that the HAM, ViT, and GEM modules significantly enhance performance. This finding underscores these modules’ ability to capture cellular morphology information in tissue images, optimize feature representation at each latent stage, and capture higher-order associations between features across multiple latent stages.

**Figure 4.**
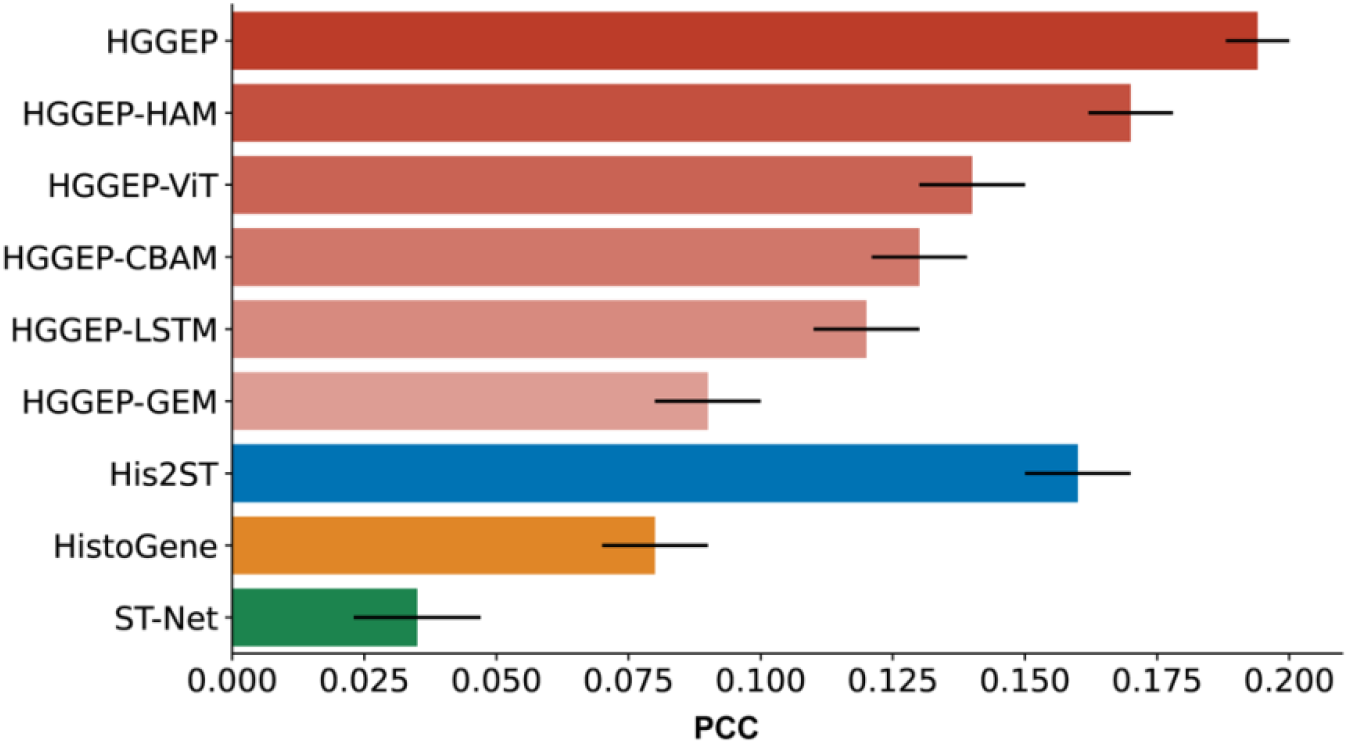
Ablation studies of our HGGEP model. As the number of modules removes, the mean PCC of the model gradually declines, underscoring the substantial contribution each module makes to the overall performance of the HGGEP model.

### Evaluation of predicted gene expression

Next we have compared the predicted gene expression between the HGGEP model, Hist2ST, and HisToGene. As shown in Figure S4, most dots are positioned above the diagonal line, indicating that the HGGEP model achieves superior gene expression predictions. It is noteworthy that the top genes identified in the HER2+ dataset (*GNAS, UBA5*2, *MUCL1*) and the cSCC dataset (*EIF5, MSMO1, STMN1*) indicate significant biological relevance. In the HER2+ tumor microenvironment, *GNAS* is implicated with critical roles in cell signal transduction^34^, potentially influencing the biological processes including the proliferation and metastasis of tumor cells^35^. *UBA5*2, closely associated with ubiquitin-protein conjugation^36^, may significantly impact the treatment response and prognosis of HER2+ tumors due to its aberrant expression^37^. Additionally, *MUCL1*, involved in cell adhesion and tumor microenvironment regulation, plays a crucial role in HER2+ tumors and offers potential insights for precision therapy^38^. On the other hand, in the cSCC environment, *EIF5* is implicated in protein synthesis regulation and cell proliferation^39^. *MSMO1* participates in cholesterol biosynthesis^40^, and *STMN1* may influence biological processes such as cell division and migration^41^. Therefore, accurate prediction of those genes contributes to exploring the molecular mechanisms driving cSCC development, thus gaining a better understanding of tumor cell activities in the microenvironment.

For a more intuitive comparison of top gene prediction performance across different models, we conducted the visualization of predicted genes. To ensure a fair comparison, we select four genes mentioned in Hist2ST (*GNAS, FASN, MY1*2*B*, and *SCD*) and present their predicted gene expressions by each model in **Figure 5**. This figure demonstrates our model achieves the best prediction performance. For example, our model achieves better prediction of the *GNAS* gene expression (PCC= 0.637), compared with the competitors (Hist2ST, PCC = 0.591; HisToGene, PCC = 0.493). Similarly, for the genes *FASN, MY1*2*B*, and *SCD*, our model demonstrates consistently better prediction performance. These results support that our model not only excels in overall performance but also demonstrates outstanding performance in predicting specific genes with biological significance.

**Figure 5.**
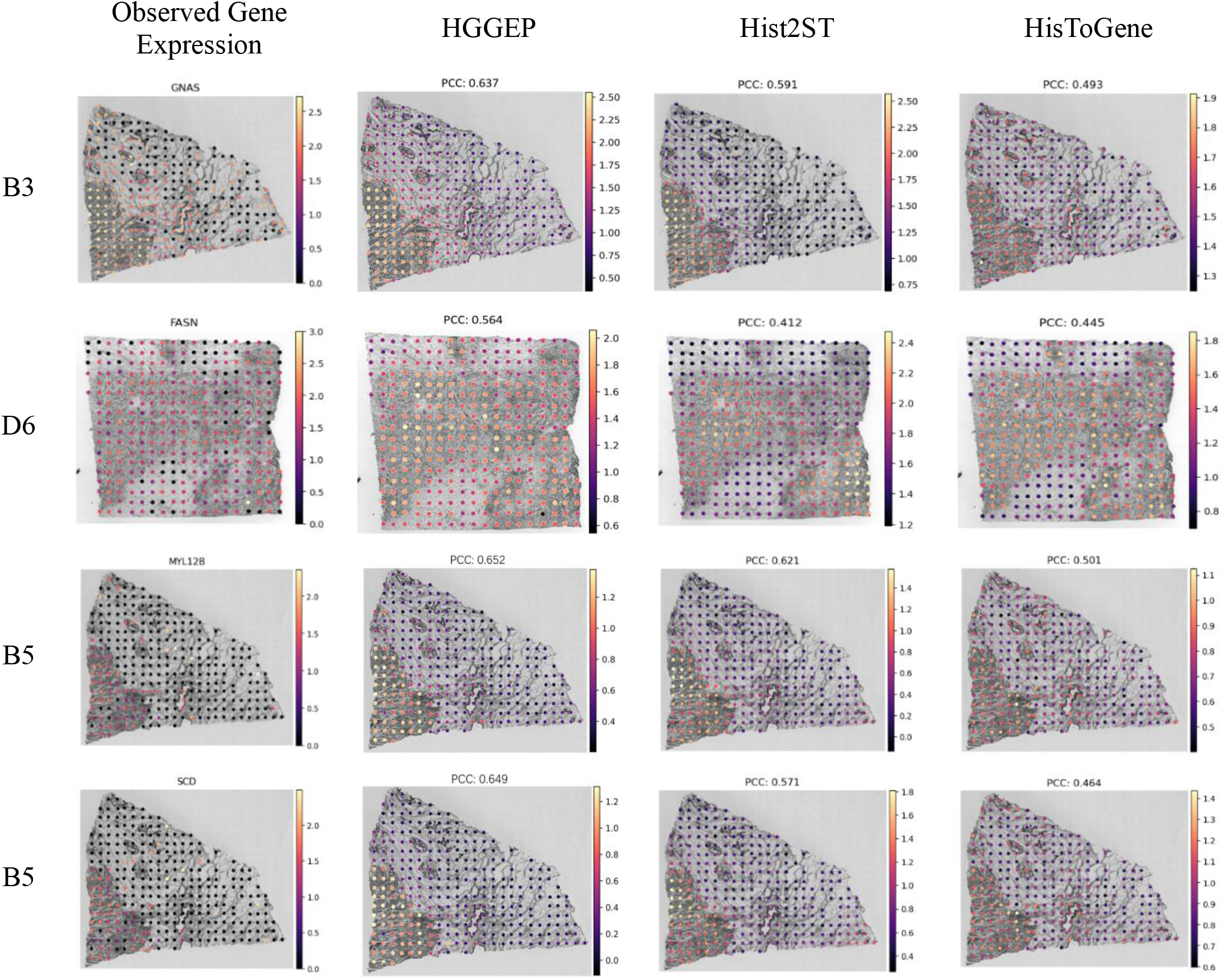
Visualization of predicted genes. The top predicted genes across all tissue sections by HisToGene in the HER2+ dataset, where the p-value for each tissue section was obtained in the association test between the predicted and observed gene expression.

### Evaluation of spatial region detection

To further demonstrate the accurate gene expressions predicted by our model, we evaluate the spatial region detection capability across the entire histology image accordingly. We performed Leiden clustering analysis on the predicted gene expressions of each method, as illustrated in **Figure 6**. Here we utilized six sections (B1, C1, D1, E1, F1, and G2) of the HER2+ dataset, with available annotations from expert professionals. As depicted in **Figure 6**, our model consistently outperforms the latest Hist2ST and HisToGene models, achieving nearly optimal detection performance. Across the six sections, our model surpasses the runner-up model Hist2ST by approximately 11% in average performance. Particularly noteworthy is the outstanding performance in section C1, where our model significantly improves the ARI index by 30% compared to Hist2ST, demonstrating the superior precision of our HGGEP model in region detection.

**Figure 6.**
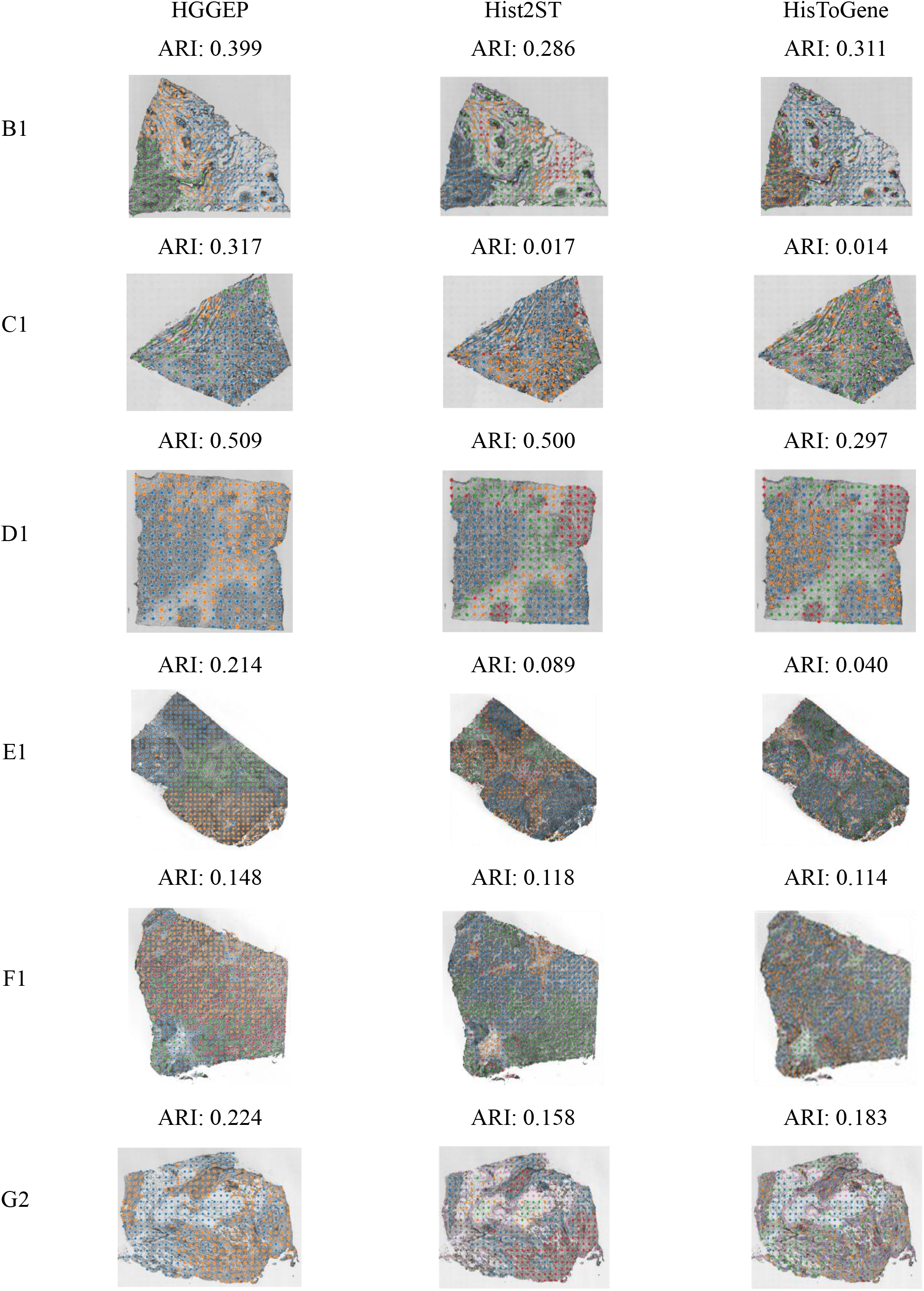
Spatial domain detection based on predicted gene expressions. The first column presents the observed gene expression clustering results, while the last three columns show the clustering outcomes for gene expression as predicted by different methods (HGGEP, Hist2ST and HisToGene).

## DISCUSSION

Gene expression prediction is a pivotal focus in current research, with its extensive applications making it a central theme in scientific exploration. However, many existing methods face limitations due to their reliance on costly data, which constrains further research progress. In response to this challenge, some researchers have turned to a more cost-effective strategy, utilizing histology images to unveil gene expression. In this work, we have introduced HGGEP, a cutting-edge hypergraph neural network model tailored for the precise prediction of gene expression from histology images. HGGEP has been applied to a total of 44 sections, showcasing robust performance in predicting gene expression and underscoring the pivotal role of histology images in this process.

Our HGGEP model outperforms existing methods in three aspects. First, we enhance the model’s ability to perceive morphological information through the gradient enhancement module, thus enabling the model to capture the relationship between cell morphology and gene expression. Second, features at different stages in the image have different information, usually high-level features have more semantic information, while low-level features carry more detailed features. Therefore, we use a lightweight backbone to extract different levels of information and then use this attention mechanism to optimize the features at each stage. Lastly, we fuse features from various stages, establishing global associations among spots to achieve comprehensive information integration.

Though HGGEP presents superior performance, we anticipate key areas for future exploration and improvement. First, HGGEP still relies on spot-level predictions for the gene expression of corresponding images and does not explore the strong correlation between individual cell information and gene expression within the images. In the future, we will focus on predicting gene expression based on cell-level or even pixel-level data. Second, most existing methods rely on large amounts of labeled data for learning, thus limiting the ability of the model to generalize to small sample domains. To overcome this limitation, future work could consider fine-tuning based on large vision models in the biomedical field^42, 43^. By leveraging the wealth of knowledge that already exists in large vision models, we can enhance the generalizability of the models to a wider range of application scenarios. Third, existing methods typically use histology images as input for predicting gene expression. Future research will consider introducing more modal information as input to help establish mapping relationships between images and gene expression, such as molecular biology data or pathology data. By integrating multimodal information and leveraging advanced hypergraph neural network methods for modeling, we aim to enhance the accuracy and comprehensiveness of gene expression prediction. In conclusion, the experiments in this paper demonstrate the power of HGGEP in predicting gene expression based on histology images. Moreover, future work could focus on enhancing model generalization capabilities, including the use of large vision models and multimodal information, to better accommodate the diversity of biomedical data and scenarios.

## MATERIALS AND METHODS

### Data processing

To validate the efficacy of our proposed method, we utilized the spatial transcriptomics datasets comprising histology images and gene expression data at spot locations. We primarily leveraged the HER2+ and cSCC datasets profiled from 32 and 12 tissue slides, respectively, with a total of 9,612 spots and 6,630 spots respectively. The tissue images in the Her2+ and cSCC datasets correspond to 785 and 171 highly variable genes, respectively. For each histology image in the datasets, we segment it into multiple sub-images based on the positions of spots. Each sub-image is cropped by 112×112 pixels around the spot’s center. The input sub-image features are annotated as *x*_*in*_ ∈ *R*^N×3×112×112^, where N is the number of spots within the histology image.

During the validation phase, we adopt a consistent leave-one-out cross-validation strategy. Taking the HER2+ dataset as an example, for each section, we train the model on the remaining 31 sections and validate it on that leave-out section.

### HGGEP model

In contrast to prior methods, HGGEP significantly improves the prediction accuracy of gene expression from histology images. The detailed structure of HGGEP model is illustrated below.

### Gradient enhancement module

To enrich the cell morphological information closely associated with gene expression, this paper proposes the GEM, which enhances the model’s perception of cell morphology through difference convolution^44, 45^. This module comprises two key components: 1) the convolution process that convolves the sub-images; 2) the gradient enhancement process that enhances the cell morphology information by difference convolution.

**Figure 1** illustrates the implementation of GEM with the steps of convolution and difference operations. Specifically, a 3 × 3 convolutional kernel is applied to a 5 × 5 input split histology image, resulting in a downsampled feature map of size 3 × 3. Subsequently, a gradient enhancement operation is performed on the resulting feature map, aiming to utilize the differences between neighboring pixels in the feature map to enhance its gradient information. Traditional convolution process can be expressed as:

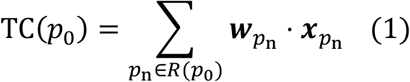

where *p*_0_ represents the central position of the local receptive field *R(p*_0_), *p*_*n*_ denotes the relative position of other pixels within the receptive field *R(p*_0_) = {(−1, −1), (−1,0), …, (0,1), (1,1)}, 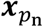 denotes the pixel value at position *p*_*n*_ in the input feature map, 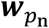 is a learnable parameter. To achieve gradient enhancement, we deform the traditional convolution TC*(p*_0_) to difference convolution DC*(p*_0_):

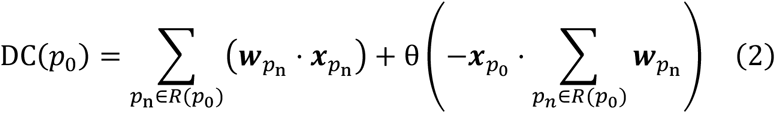

where *θ* is a hyperparameter that regulates the balance between semantic and gradient information, and we set it to 0.7 default. When *θ* = 0, the difference convolution is the same as the traditional convolution operation. Given the input sub-images ***x***_*in*_ ∈ ℝ^*N*×3×112×112^, each of the convolution (*F*_*TC*_and *F*_*DC*_) exports the *F*_*TC*_ *ϵ* ℝ^*N*×6×56×56^ and *F*_*DC*_ *ϵ* ℝ^*N*×6×56×56^. The latent features from GEM module are *z*^*GEM*^ ∈ ℝ^*N*×3×56×56^. The entire GEM operates as follows:

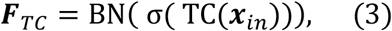

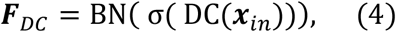

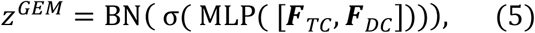

where *z*^*GEM*^ is the enhanced latent feature map, *σ* is ReLU activation function,BN is Batch Norm. Here the Multi-Layer Perceptron (MLP) layer is used to adjust the number of feature channels, which then serve as input into the subsequent backbone network.

### Backbone and multiple latent stage feature

With the GEM-enhanced feature map *z*^*GEM*^, the backbone network, CBAM^25^, and ViT^26^ are employed to extract and optimize multiple latent stage feature information from the *z*^*GEM*^, enhance the model’s prowess in latent feature modeling, and capture global relationships among spots at each latent stage.

For the backbone network, we chose a lightweight shufflenet V2^24^ to optimize computing efficiency. This network produces output across five stages, with diminishing latent feature map sizes as the network deepens. For subsequent processing, we leverage features from the last three stages from the backbone, feeding them into the CBAM. As shown in **Figure 1**, CBAM utilizes both Channel Attention (CA) and Spatial Attention (SA) for adjusting the weight of each channel/spot by considering global information. With these attention mechanisms, CBAM enhances the network’s comprehension of vital features in the image and improves the model’s performance and generalization capabilities.

Specifically, the GEM output features *z*^*GEM*^ ∈ ℝ^*N*×3×56×56^ are first fed into the shufflenet V2 backbone network to get the embeddings at 5 latent stages [***S***_1_ ∈ ℝ^*N*×24×29×29^, ***S***_2_∈ℝ^*N*×48×15×15^, ***S***_3_ ∈ ℝ^*N*×96×8×8^, ***S***_4_ ∈ ℝ^*N*×192×4×4^, ***S***_5_ ∈ ℝ^*N*×1024×4×4^]. The embeddings of the last 3 latent stages are selected for subsequent analysis. Subsequently, we input these embedding features into the CBAM to optimize the features of each latent stage separately. Specifically, for ***S***_*i*_, *i* ∈ {3,4,5}, the CBAM module performs as below:

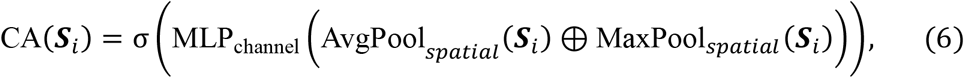

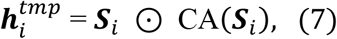

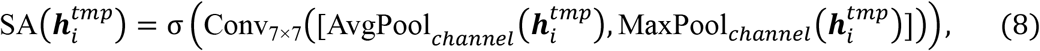

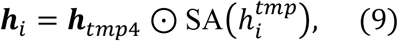

where the symbol *⊕* denotes element-wise addition, ⊙ represents element-wise multiplication, and *σ* is the ReLU activation function. AvgPool_*spatial*_ (***S***_*i*_) a nd MaxPool_*spatial*_ (***S***_*i*_) de note the average and maximum pooling operations on the embedding feature ***S***_*i*_, respectively, and the output of both is ℝ^*N*×192×1×1^. As shown in equation (6), the outputs of the two pooling operations are element-wise added. They subsequently enter the MLP_channel_ for channel scaling, with the scaling factor set to 16 by default (equation 6). For spatial attention SA 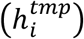, AvgPool_*channel*_ 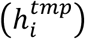 and MaxPool_*channel*_ 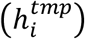 perform average and maximum pooling for the channels, respectively, followed by the 7 × 7 convolution. It is worth noting that CBAM-optimized hidden features maintain their original input dimension, i.e., the hidden features ***h***_*i*_ after CBAM optimization are ***h***_3_ ∈ ℝ^*N*×96×8×8^, ***h***_4_∈ℝ^*N*×192×4×4^, ***h***_5_ ∈ ℝ^*N*×1024×4×4^.

To facilitate subsequent processing, we uniformly adjust the above three hidden features [***h***_3_, ***h***_**4**_, ***h***_**5**_] to ℝ^*N*×1024^ by linear and dimension transformation. These hidden features are fed into the ViT module separately and its self-attention mechanism is utilized to capture the associations among the spots in WSI. For ***h***_*i*_, *i* ∈ {3,4,5}, the ViT module is implemented as follows:

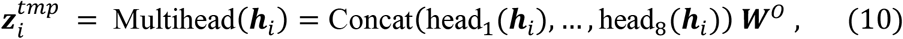

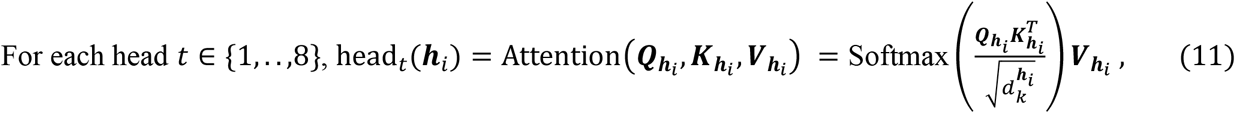

where 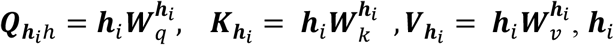, represents one of the input features 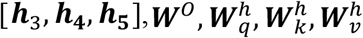denote the learnable weight matrices, 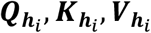 are the matrices of queries, keys, and values obtained by linear transformation of ***h***_*i*_, respectively. The parameter 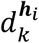 is used to scale the denominator in the dot-product attention, controlling the scaling of attention weights. These operations enable the model to dynamically model the associations among spots, leading to an optimal allocation of attention. After the optimization of the multi-head attention mechanism, the dimension of the ViT output 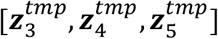 remain constant at ℝ^*N*×1024^.

Finally, a Feed-Forward Network (FFN) is applied for additional non-linear transformations and representation learning of the hidden features from each stage:

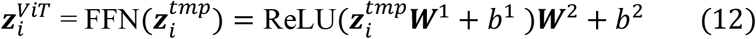

where ***W***^*i*^ and *b*^*i*^ are the weight matrix and bias vector, respectively. After ViT optimization, it is obtained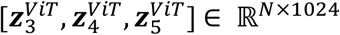.

### Hypergraph association module

The feature representations of spots at each latent stage are derived by encoding histology images through the model’s front-end image encoder, which encompasses the GEM, Backbone, CBAM, and ViT modules. However, the above modules are limited to treating the features of each stage independently and do not explore the association among the multiple latent stage features 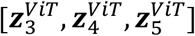. Herein, we first fuse the multiple latent feature maps described above, and then introduce a Hypergraph Association Module (HAM) aimed at capturing high-order association among spots. To comprehensively model feature associations at varying distances, we adopt a global modeling approach based on Euclidean distance and a local modeling approach based on nearby positions.

Specifically, with the summed element by element of 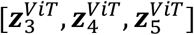, ***H***_*in*_ *ϵ* ℝ^*N*×1024^ is obtained and fed into the HAM. We use ***v***_*i*_ as the node and the attributes of the node ***v***_*i*_ is ***m***_*i*_ *ϵ* ℝ^1×1024^. To establish hyperedges among nodes, we propose a method that combines Euclidean distance and nearby positions metrics to generate an incidence matrix. Initially, we measure the Euclidean distance between the current node ***v***_*i*_ and other nodes. This effectively models relationships between distant nodes ***v***_*i*_ and ***v***_*j*_:

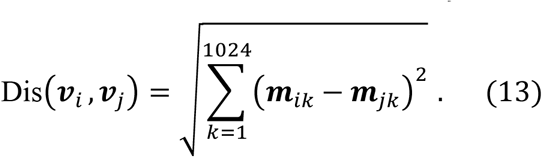

Simultaneously, considering that neighboring nodes on the coordinates usually have similar genetic phenotypes, we also build hyperedges based on the positional relationship between nodes. Assume that the coordinates of node ***v***_*i*_ are (*x*_*i*_, *y*_*i*_) and the coordinates of node ***v***_*j*_ are (*x*_*j*_, *y*_*j*_), the positional relationship is encoded as:

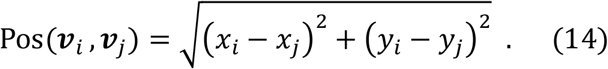

By combining the two aforementioned metrics, we establish the relationship between the current node and other nodes. To ensure a balanced contribution of the two metrics to the final incidence matrix, normalization method is applied:

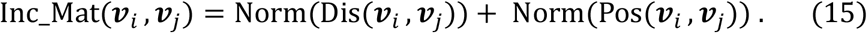

Next, we construct the hyperedges based on the incidence matrix. For each node, we find the top K nearest nodes based on the incidence matrix, and then construct a hyperedge from these K nodes. The hyperedges, along with their attributes and features ***H***_*in*_ *ϵ* ℝ^*N*×1024^, are input into a hypergraph convolutional network. Throughout the convolution process, the model enhances its understanding of inter-node dependencies, encompassing relationships among nodes as well as between nodes and hyperedges. For the concrete implementation of hypergraph convolution^28, 29^, this study leverages the HypergraphConv module from the torch_geometric library:

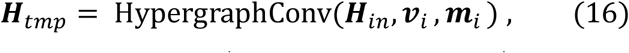

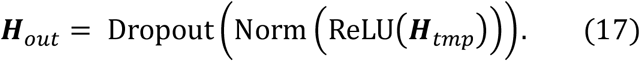

The output feature obtained after the HAM module is ***H***_*out*_ *ϵ* ℝ^*N*×1024^.

Finally, to obtain the final gene prediction, a Long Short-Term Memory (LSTM) module and MLP module are introduced in this paper. LSTM models the association among features by treating the input multiple latent stage features as time series. The features of the four stages 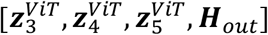 are concatenated together to obtain ***L***_***in***_ *ϵ* ℝ^4×*N*×1024^, which is used as an input to the LSTM. Subsequently, the first dimension of the LSTM output features is average pooled to obtain the average vector ***L***_***out***_*ϵ* ℝ^*N*×1024^. Finally, MLP is used to map the latent feature ***L***_***out***_ *ϵ* ℝ^*N*×1024^ to the number of final predicted genes, which is ℝ^*N*×785^ for HER2+ and ℝ^*N*×171^ for cSCC dataset.

### Model hyperparameter configuration

The hyperparameters used in the HGGEP model are listed specifically. (1) GEM: the parameter θ in Equation 2 is set to 0.7 and the feature channel variations progress from 3→6→3. (2) The backbone network: we utilize the shufflenet_v2_x0_5 model, pre-trained on the ImageNet dataset, to extract features from the final three stages for subsequent modules. (3) CBAM: the scaling factor in the channel attention in Equation 6 is configured to shrink by 16 x. Subsequently, for ease of subsequent processing, we reduce these features from the last three stages to ℝ^n×1024^, where n represents the number of spots. (4) ViT: the whole ViT module is iterated four times. The number of attention heads is set to 8, and the FeedForward component includes 2 MLP layers. (5) HAM: when constructing the incidence matrix, a fixed number of adjacent nodes is set to 3, and nodes retain their own indices. The convolution process involves 2 layers of hypergraph convolution, with the hidden layer being half the size of the input. (6) LSTM: This encompasses input and hidden state dimensions of 1024 each, along with a layer count of 4. (7) MLP: After output normalization, a single MLP layer is directly employed to predict the final gene output.

### Loss function

Mean Squared Error (MSE) loss is utilized to measure the average squared difference between the predicted gene expression values and the ground truth for each gene. Meanwhile, given the prevalent zeros in the spatial transcriptomics data, we also include the Zero-Inflated Negative Binomial (ZINB ^46^) loss. This loss is rooted in the zero-inflated negative binomial distribution, amalgamating the negative binomial distribution with a zero-inflation component. The total loss of the HGGEP model is a fusion of MSE and ZINB loss, for the task of gene prediction from histology images.

### Evaluation metrics

To comprehensively evaluate the model’s performance, this paper utilizes three metrics: Pearson Correlation Coefficient (PCC), Structural Similarity Index Measure (SSIM), and Root Mean Square Error (RMSE).

Pearson Correlation Coefficient (PCC) is used to assess the performance of benchmarking models. This coefficient, a statistical measure for assessing the linear relationship between two continuous variables, is widely employed to quantify the strength and direction of the linear correlation between these variables. The PCC ranges from -1 to 1, with the progression from -1 to 1 indicating a shift from negative to positive correlation. The specific calculation method is as follows:

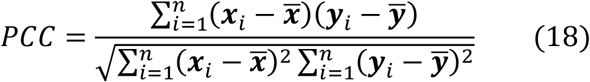

where 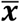 and 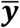 represent the means of variables ***x*** and ***y***, respectively, and *x*_*i*_ and ***y***_*i*_ denote the *i*_*th*_ observations.

Structural Similarity Index Measure (SSIM) is a perceptual metric that quantifies data quality degradation caused by processing such as data compression or transmission. Unlike traditional metrics like Mean Squared Error (MSE), SSIM considers changes in structural information, luminance, and contrast. SSIM values range from -1 to 1, where a value closer to 1 indicates that the two images are highly similar. The SSIM calculation involves three components: luminance (l), contrast (c), and structure (s). The formula is given by:

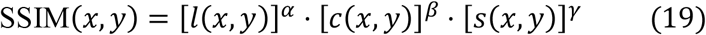

Typically, the values for *α, β*, and *γ* are set to 1, leading to the simplified formula:

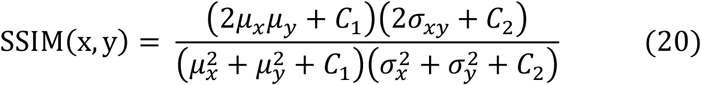

where *μ*_*x*_ and *μ*_*y*_ are the mean intensities of images *x* and *y*, 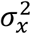 and 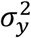 are the variances, *σ*_*xy*_ is the covariance, and *C*_1_ and *C*_2_ are small constants to stabilize the division.

Root Mean Square Error (RMSE) is used to evaluate the performance of benchmarking models by measuring the differences between predicted and actual values. RMSE is a widely-used statistical metric that quantifies the magnitude of prediction errors. The RMSE value is always non-negative, with a lower RMSE indicating better model performance. The RMSE is calculated as follows:

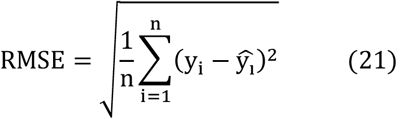

Where *y*_*i*_ represents the actual observed values 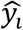 represents the predicted values, and *n* is the number of observations. The formula takes the square root of the average of the squared differences between the predicted and actual values, providing a measure of the model’s predictive accuracy.

These metrics, PCC, SSIM, and RMSE, together provide a comprehensive evaluation of the model’s performance, capturing different aspects of prediction accuracy and data quality.

### Training parameters

During training, the model utilizes a learning rate of 0.00001, runs for a maximum of 400 epochs, and undergoes testing every 5 epochs. The experiments are conducted on an Ubuntu 20.04 system equipped with 128GB of RAM and an A6000 GPU featuring 48GB of memory.

## CODE AVAILABILITY

All source codes and trained models in our experiments have been deposited at https://github.com/QSong-github/HGGEP.

## DATA AVAILABILITY

The spatial transcriptomics datasets used in this study include the (1) HER2-positive breast tumor ST datasets, which are available at https://github.com/almaan/her2st/; (2) 10x Visium data of human cutaneous squamous cell carcinoma are publicly available in the Gene Expression Omnibus (GEO) (GSE144240).

## MATERIALS & CORRESPONDENCE

Correspondence and requests for materials should be addressed to QS or YZ.

